# Network-Guided Discovery of Influenza Virus Replication Host Factors

**DOI:** 10.1101/344861

**Authors:** Emily E. Ackerman, Eiryo Kawakami, Manami Katoh, Tokiko Watanabe, Shinji Watanabe, Yuriko Tomita, Tiago J. Lopes, Yukiko Matsuoka, Hiroaki Kitano, Jason E. Shoemaker, Yoshihiro Kawaoka

**Affiliations:** Department of Chemical & Petroleum Engineering, Swanson School of Engineering, University of Pittsburgh, Pittsburgh, PA, 15213, USA; Division of Virology, Department of Microbiology and Immunology, Institute of Medical Science, University of Tokyo, Tokyo 108-8639, Japan; ERATO Infection-Induced Host Responses Project, Japan Science and Technology Agency, Saitama 332-0012, Japan; The Systems Biology Institute, Minato-ku, Tokyo 108-0071, Japan; Department of Pathobiological Sciences, School of Veterinary Medicine, University of Wisconsin-Madison, 575 Science Drive, Madison, WI 53711, USA; Laboratory for Disease Systems Modeling, RIKEN Center for Integrative Medical Sciences, 1-7-22 Suehiro, Tsurumi, Yokohama, Kanagawa 230-0045, Japan; Okinawa Institute of Science and Technology, Onna-son, Okinawa 904-0495, Japan; Department of Computational and Systems Biology, School of Medicine, University of Pittsburgh, Pittsburgh, PA, 15213, USA; Department of Special Pathogens, International Research Center for Infectious Diseases, Institute of Medical Science, University of Tokyo, Minato-ku, Tokyo 108-8639, Japan

## Abstract

The position of host factors required for viral replication within a human protein-protein interaction (PPI) network can be exploited to identify drug targets that are robust to drug-mediated selective pressure. Host factors can physically interact with viral proteins, be a component of pathways regulated by viruses (where proteins themselves do not interact with viral proteins) or be required for viral replication but unregulated by viruses. Here, we demonstrate a method of combining a human PPI network with virus-host protein interaction data to improve antiviral drug discovery for influenza viruses by identifying target host proteins. Network analysis shows that influenza virus proteins physically interact with host proteins in network positions significant for information flow. We have isolated a subnetwork of the human PPI network which connects virus-interacting host proteins to host factors that are important for influenza virus replication without physically interacting with viral proteins. The subnetwork is enriched for signaling and immune processes. Selecting proteins based on network topology within the subnetwork, we performed an siRNA screen to determine if the subnetwork was enriched for virus replication host factors and if network position within the subnetwork offers an advantage in prioritization of drug targets to control influenza virus replication. We found that the subnetwork is highly enriched for target host proteins – more so than the set of host factors that physically interact with viral proteins. Our findings demonstrate that network positions are a powerful predictor to guide antiviral drug candidate prioritization.

**IMPORTANCE:** Integrating virus-host interactions with host protein-protein interactions, we have created a method using these established network practices to identify host factors (i.e. proteins) that are likely candidates for antiviral drug targeting. We demonstrate that interaction cascades between host proteins that directly interact with viral proteins and host factors that are important to influenza replication are enriched for signaling and immune processes. Additionally, we show that host proteins that interact with viral proteins are in network locations of power. Finally, we demonstrate a new network methodology to predict novel host factors and validate predictions with an siRNA screen. Our results show that integrating virus-host proteins interactions is useful in the identification of antiviral drug target candidates.

## INTRODUCTION

Viruses such as influenza virus hijack and reprogram host cellular machinery to perform virus replication tasks. Influenza outbreaks have a major impact on public health and the global economy each year(1, 2). While annual vaccinations provide some protection, vaccination effectiveness is impaired by antigenic drift and availability issues(3, 4). Recent sporadic human infections with avian viruses of H5N1 and H7N9 subtypes have raised concerns about the pandemic potential of these viruses(5–8). Antiviral drugs that target influenza viral proteins are available(9, 10) but drug resistant strains have emerged(11, 12). Therefore, there is an urgent need to identify drug targets that are robust to virus mutation and drug-mediated selective pressure.

Understanding virus-host interactions in the context of the human protein-protein interaction (PPI) network will provide a global perspective into how influenza virus manipulates host proteins and aid in identifying host factors involved in influenza virus replication for drug targeting(13–15). The virus-host interactome is visualized in Fig. 1A. Within a PPI network, a protein’s global significance can be assessed by the protein’s network centrality: the identification of important components based on information flow across the network. Common measures include a proteins degree (number of binding partners) and betweenness (the degree to which the protein is a bottleneck in the network) though several others exist(16, 17). Network centrality has emerged as a valuable tool for studying proteins associated with cancer(18, 19) and drug targeting(19–22). PPI network-based approaches have recently been utilized in influenza virus studies to identify and study potential factors involved in virus replication(23–27). Network studies have demonstrated that virus interacting host proteins tend to have a high network significance based on a variety of network metrics (including betweenness and degree) for several viruses including influenza viruses(28) and hepatitis C virus(29). A comparative analysis of influenza virus protein and host protein interactomes has identified novel host factors that are common across the protein interactomes(30). Furthermore, meta-analysis studies have developed statistical methods to better identify host factors by leveraging data from several virus replication screens (31). Yet, a remaining question is how effectively can virus-host protein interaction data and network topology be leveraged to improve host factor identification (i.e. antiviral drug target identification).

**Figure 1.**
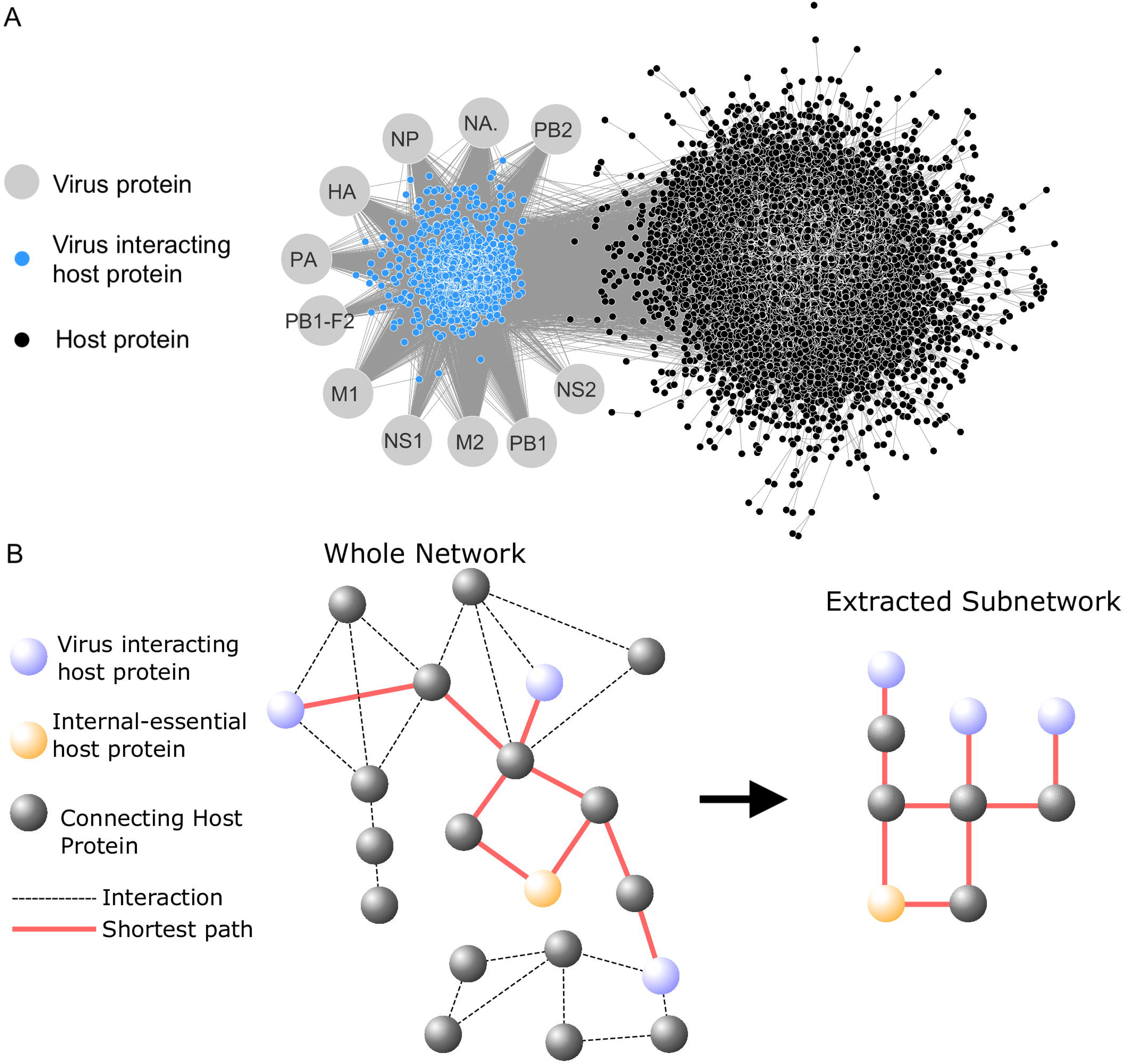
The virus-interacting network and the virus subnetwork. (a) The virus-interacting network is created from human host-PPI data combined with virus-host protein interaction data. (b) The virus subnetwork was isolated from the complete human PPI network by collecting all interactions involved in the shortest paths (red) that connect influenza virus-interacting proteins (blue) to human proteins essential to virus replication (e.g. the internal-essential proteins; colored orange). The connecting proteins (colored black) are candidates to be evaluated for their antiviral properties.

Here, we demonstrate a method of integrating virus-host protein interaction data into a human PPI network to prioritize host proteins as antiviral drug target candidates. First, we analyzed a set of 1,292 human proteins identified previously as having interactions with influenza virus proteins(32), 299 of which were found to significantly inhibit influenza virus replication during an siRNA virus replication screen (Fig. 1A). Consistent with previous studies, we show that virus-interacting human proteins tend to be in positions essential to PPI network information flow and are closely clustered within the PPI network. We then isolated the subnetwork of the human PPI network that connects virus-interacting host proteins to non-interacting, host factors (referred to as “internal”) that were identified to be important for influenza virus replication in a study and re-evaluated in this work (33) (Fig. 1B). Candidate proteins connecting virus-interacting host proteins to internal host factors were selected based on their betweenness within this subnetwork and evaluated by viral replication screen. Betweenness was selected under the hypothesis that selecting network bottlenecks (i.e. high betweenness proteins) would limit the opportunity for the virus to engage host machinery through alternative pathways. The fraction of proteins tested which significantly reduced virus replication (i.e. the hit rate) was compared to the hit rate observed in a genome-wide screen, the hit rate when screening virus-interacting proteins (the virus’ nearest neighbors in the network) and the hit rate observed when screening host factor identified in a previous study(33).

## RESULTS

### Host proteins that interact with influenza virus proteins are central to the PPI network

Studies have shown that proteins in network positions that are essential for information flow within a PPI network (e.g. high degree or high betweenness) are more likely to be associated with diseases(34, 35) or drugs with known, dangerous side-effects(19, 36). Using a human PPI network, we analyzed the network topology characteristics of virus-interacting and non-virus-interacting host proteins. In a previous study, we identified 1,292 host proteins that co-precipitated with at least one of 11 influenza virus proteins (viral PB2, PB1, PA, HA, NP, NA, M1, M2, NS1, NS2, and PB1-F2 proteins)(32). These proteins are referred to as “virus-interacting proteins”. We mapped the interaction data onto a human PPI network developed from the Human Integrated Protein-Protein Interaction rEference (HIPPIE) database(37). After constraining the network to highly confident interactions (see Methods), the PPI consisted of one large network (9,969 proteins and 57,615 interactions) which contained 1,213 influenza virus-interacting host proteins and 86 smaller networks that contained 7 or fewer proteins (the majority only containing 2 proteins) and no influenza virus-interacting proteins. The smaller networks were removed from further consideration.

Virus proteins were significantly more likely to interact with host proteins that were in positions of high regulatory importance in the human PPI network. For every protein, the degree (number of neighbor proteins) and betweenness(38) (measure of the shortest paths between all other proteins in the network that include the protein in question) were calculated. On average, the degree of virus-interacting host proteins was twice the degree of all proteins in the network (Fig. 2A; the median degree of virus-interacting proteins = 10; the median degree of all proteins in the network = 5; Student t-test of the log-scaled data p < 10^−16^). Virus-interacting proteins also had a significantly higher betweenness (Fig. 2B; virus-interacting proteins median betweenness = 1625.1; the median betweenness of all protein in the network = 32.8; Mann-Whitney U test of the log-scaled data p < 10^−16^). We also compared to the median betweenness when removing proteins with a betweenness of zero. Virus-interacting proteins still had a significantly higher betweenness but the population medians were closer in value (virus-interacting proteins median betweenness = 3981.1; the median betweenness of all protein in the network = 1584.8; Mann-Whitney U test of the log-scaled data p = 8.2*10^−16^). The tendency for virus proteins to bind host proteins that had a higher degree and betweenness was consistent when analyzing the interaction partners of each virus protein separately (Fig. S1; pairwise t-test of the log-scaled data. All p < 0.01 except for betweenness of NS2-interacting proteins which was not significantly distinct from the betweenness of the full PPI). This indicates that influenza virus proteins selectively interact with human proteins that can strongly regulate cellular behavior. These results are consistent with literature findings for HCV and Dengue virus(39, 40) and with a previous study which used a yeast two-hybrid approach to identify influenza virus interacting host proteins for 10 of the 11 virus proteins (28). Further, these are characteristics that generalize to each virus protein’s interacting partner; suggesting that all 11 virus proteins have a role in manipulating cellular machinery.

**Figure 2.**
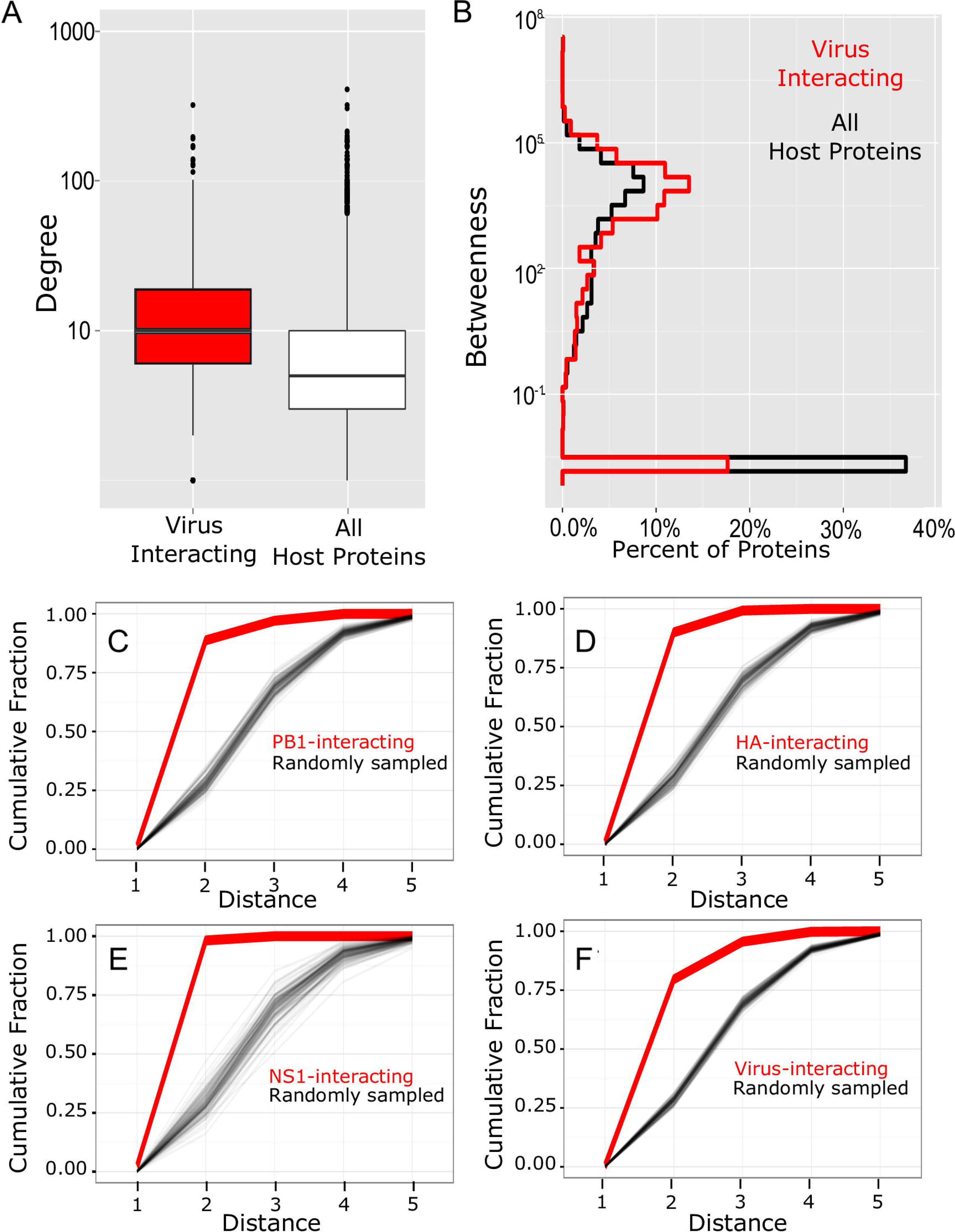
The network topological characteristics of virus-interacting host proteins. The distributions of the (A) degree and (B) betweenness of virus-interacting proteins and all proteins in the human PPI network. An ε = 0.01 was added to the betweenness to facilitate log scaling. The cumulative distributions (thick, red lines) of the shortest distances connecting host proteins in the PPI network that interact with viral (C) PB1, (D) HA, (E) NS1 proteins or (F) the set of all viral proteins. As a control, the cumulative distribution of distances was iteratively determined (N=100) by randomly sampled host proteins in the PPI network (thin, black lines). The number of proteins sampled on each iteration was equal to the number of interacting host proteins of each virus protein (or set of viral proteins).

### Influenza virus-interacting host proteins are closely connected in the human PPI network

Next, we evaluated if virus-interacting proteins tend to cluster closely to one another in the PPI network. A previous study suggested that host factors of viral replication are closely clustered within the network but did not assess the topological characteristics of virus-interacting host proteins (41). Functionally related proteins are often observed to be closely clustered in PPI networks(42, 43). Knowing that influenza virus proteins manipulate multiple host cell functions to promote replication, these previous studies suggest that the interaction partners of viral proteins should be closely clustered by host function. If true, neighboring cluster proteins could serve as possible alternatives for influenza virus to manipulate each host function.

We quantified how close each virus proteins’ interacting host proteins are within the network by calculating the shortest distances required to connect all of the host proteins that interact with a viral protein, creating a distribution of distances. The cumulative distribution details the fraction of host proteins that could be connected to other host proteins that bind the same viral protein in *n* or fewer steps. As a control, we determined the cumulative distribution of distances that result from randomly sampled proteins in the network. For a single iteration, we created a set of random proteins. The size of the set was determined by the number of proteins which interact with the virus protein of interest (e.g. PB1 has 322 interacting host proteins, therefore 322 proteins were randomly selected from the network; Fig. 2C-F). The distributions of distances connecting all of the randomly sampled proteins was calculated. This was process was repeated 100 times.

We found that virus-interacting host proteins are very significantly clustered within the PPI network. The set of proteins that interact with a viral protein are significantly more closely clustered in the network than expected by chance (Fig. 2C-F, p < 0.01 when comparing the median distance of the virus-interacting proteins to the median distance of randomly sampled proteins). Generally, ~25% of the randomly sampled proteins are connected by 2 or fewer interactions while 88.7% of PB1-interacting proteins, 90.0% of HA-interacting proteins, 98.2% of NS1-interacting proteins, and 79.6% of all host proteins that interact any influenza virus protein are connected by 2 or fewer interactions. Collectively, these results support that viral proteins are selectively targeting closely clustered host proteins.

We next evaluated if influenza interacting proteins are often components of a common protein complex. To do so, we determined the fraction of all influenza virus interacting proteins pairs (735,078 pairs in total) that appear within a protein complex and compared that fraction to the fraction of all protein pairs (49,685,496 total pairs) in the PPI that appear in a protein complex. Mammalian protein complex information was downloaded from CORUM (a comprehensive resource of mammalian protein complex data)(44). We found that 1.5% of all virus interacting protein pairs are involved in a complex where as only 0.066% of all proteins pairs in the PPI are involved in a complex. In sum, influenza virus proteins are closely clustered and 22.4 times more likely to be involved in a protein complex than randomly selected proteins.

### Constructing the influenza virus-host subnetwork

Network analysis of virus-interacting host proteins demonstrates that viral proteins preferentially interact with closely connected host proteins that are in positions central to information flow across the human PPI network; suggesting that it may be possible to exploit network positions to prioritize potential antiviral drug targets. We hypothesized that there exists a subnetwork consisting of pathways that *connect* virus interacting proteins to key cellular machinery that is likely to be significantly enriched for host factors. We further hypothesized that the topology of host factors within this subnetwork may provide an additional advantage in identifying host factors.

To evaluate these hypotheses, we first performed an siRNA screen of host factors identified in a previous genome-wide screen for influenza virus host factors to identify key host factors that do not interact directly with the virus (33). Poor repeatability due to differences in the experimental conditions and possibly high false negative rates (41) often characterizes siRNA screens of influenza virus replication host factors. Here, HEK293 cells were transfected with siRNAs targeting 264 non-virus interacting host factors identified in Karlas et al 2010 (two siRNAs per gene were used, as shown in Table S1; AllStars Negative Control siRNA [QIAGEN] was used as a negative control), then infected with influenza virus at 24 hours post-transfection. The culture supernatants were harvested for virus titration at 48 hours post-infection. Virus titers were determined by plaque assay. A protein was defined as a hit if the virus titers decreased by at least two log units upon transfection with an adjusted p < 0.01. Cell viability of siRNA-transfected cells was assessed using Cell-Titer Glo assay and down-regulation of mRNA levels for the hit proteins were confirmed by qRT-PCR. Of the 264 previously identified host factors tested, 71 significantly down-regulated virus replication. Of the 71, 21 were identified to directly interact with influenza virus proteins. In all, 50 of the host factors down-regulated virus growth and do not directly interact with the virus. We labeled these proteins as “internal-essential” host factors.

Next, we constructed an influenza virus specific subnetwork (process illustrated in Fig. 1B) of the shortest paths connecting virus-interacting host proteins to “internal-essential” host factors (i.e. the host factors re-verified in the siRNA screen of host factors identified in the Karlas *et al.* screen). The proteins linking internal-essential proteins to virus-interacting proteins are “connecting” candidate proteins for evaluation as host factors of virus replication. The resulting subnetwork contained 1,213 virus-interacting proteins, 38 internal-essential proteins (12 proteins were not in the PPI network), and 1,643 connecting candidate proteins (Table S2 contains the identities and centrality values for all proteins in the subnetwork). As a result of how the subnetwork is constructed, the mean degree of the virus-interacting proteins and the internal-essential proteins were lower than the mean degree of the connecting proteins (see Fig. S2A; ANOVA followed by Tukey post hoc analysis p < 0.01). While the degree of connecting proteins does not shift significantly between the total PPI network and the virus subnetwork (Fig. 3A), some proteins with low betweenness have much lower betweenness in the virus subnetwork when compared to the total PPI network (Fig. 3B). Higher betweenness nodes in the total PPI network do not demonstrate dramatic shifts in the virus subnetwork upon comparison. This shift between the total network and virus subnetwork may reveal proteins that are more or less critical to virus replication which cannot be identified in a standard PPI network analysis.

**Figure 3.**
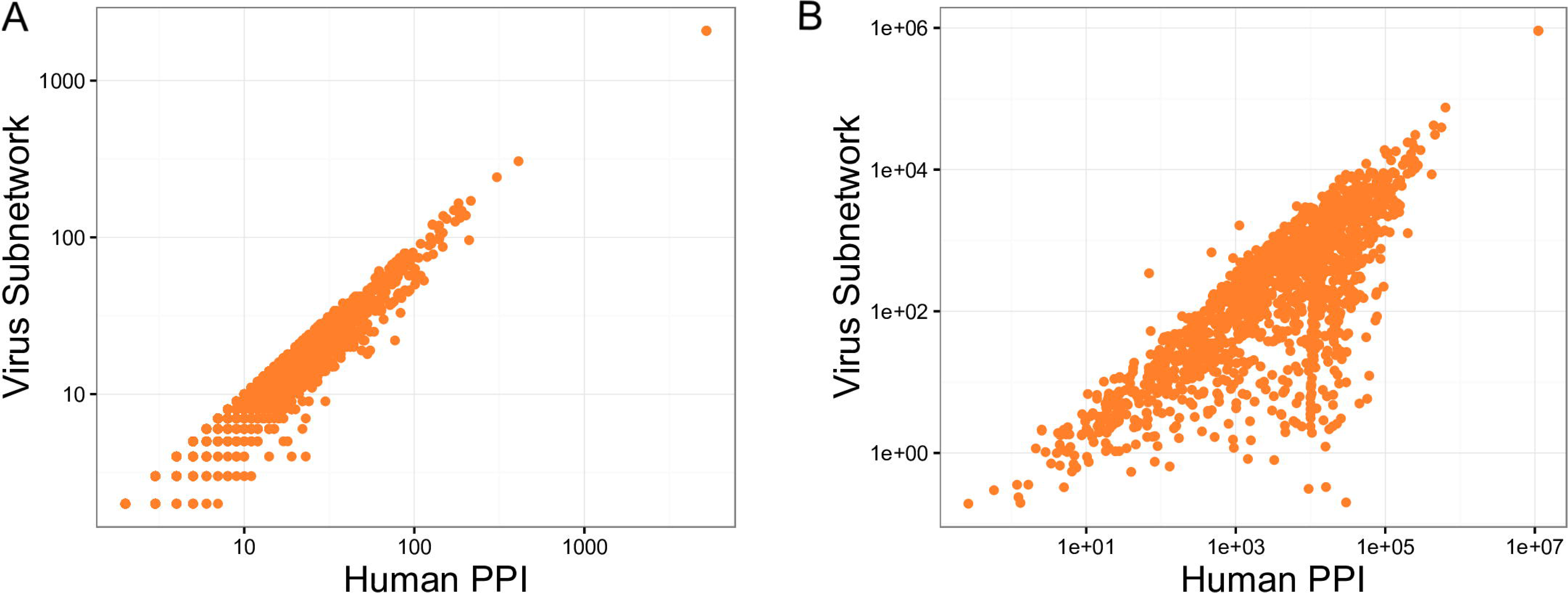
Network characteristics of the virus subnetwork. Panels (A) and (B) compare the degree and betweenness, respectively, of the connecting proteins in the whole PPI network and the virus subnetwork.

### Functional enrichment analysis of the influenza virus-host subnetwork

A functional enrichment analysis was performed using DAVID 6.8’s Functional Annotation tool(45). Analysis found that virus-interacting host proteins and connecting (non-internal-essential) proteins within the virus subnetwork are functionally distinct (see Table 1-2 for abbreviated results, see Table S3 for full results). Gene ontology and pathway analysis found that virus-interacting host proteins are primarily associated with housekeeping and viral replication processes (consistent with the results reported by Watanabe *et al.*(32)), whereas connecting proteins were associated with protein phosphorylation, histone reconfiguration and immune responses. Specifically, the immune response pathways identified are the stimulatory C-type lectin receptor signaling, T-cell receptor signaling, and Fc-epsilon receptor signaling; all of which regulate NFκB activity. Influenza virus is known to manipulate host immune response pathways (specifically NFκB regulating pathways) to promote viral replication(46, 47). These results suggest that the virus subnetwork contains functional information generally unobserved when considering virus-interacting host proteins or internal-essential proteins in isolation.

**Table 1.** Functional Enrichment Analysis of Virus Subnetwork. Functional enrichment analysis of connecting proteins within the virus subnetwork. Proteins were analyzed using DAVID.

| Cluster | Number of GO terms | Enrichment Score |
| --- | --- | --- |
| Transcription | 4 | 55.4 |
| DNA damage/repair | 3 | 19.2 |
| Protein phosphorylation | 19 | 18.7 |
| Mitosis | 5 | 18.7 |
| Histone reconfiguration | 42 | 14.4 |
| Immune response | 3 | 14.0 |
| C-type lectin receptor signaling pathway |  |  |
| T cell receptor signaling pathway |  |  |
| Zinc ion binding | 4 | 11.5 |

**Table 2.** Functional Enrichment Analysis of Virus Subnetwork. Functional enrichment analysis of virus-interacting proteins within the virus subnetwork. Proteins were analyzed using DAVID.

| Cluster | Number of GO terms | Enrichment Score |
| --- | --- | --- |
| Ribonucleoprotein/Viral transcription | 13 | 67.4 |
| Cell-cell adhesion | 3 | 46.6 |
| mRNA splicing | 9 | 41.8 |
| Nucleotide binding | 10 | 30.5 |
| Chaperone/UPR | 3 | 22.0 |
| Viral nucleocapsid | 3 | 19.0 |
| mRNA nuclear export | 4 | 17.4 |
| Nucleotide binding/ATP binding | 5 | 17.1 |
| Translation initiation factors | 11 | 13.1 |
| Proteasome/NF-kB MAPK signaling | 23 | 12.0 |

### Connecting proteins of the influenza virus-host subnetwork are more enriched for host factors than are virus-interacting proteins

To evaluate the hypothesis that the “connecting” proteins are likely to be host factors and to simultaneously evaluate if network topology can improve host factor identification, we selected 78 proteins of the subnetwork with the highest (n=39) and lowest betweenness (n=39) and conducted another *in vitro* virus replication assay. HEK293 cells were again transfected with siRNAs targeted to each of the 78 candidate protein’s genes and the procedure described previously was performed to determine the proportion of the connecting proteins tested that are host factors of influenza-virus replication. The hit rate is defined as the proportion of proteins tested that significantly down-regulated virus replication.

To evaluate the significance observed in the virus replication screen of the connecting proteins, we compared the observed hit rate to the hit rate observed in a screen the 1,292 virus-interacting host proteins in HEK293 cells (hit rate = 299/1292 = 0.23)(32), in the screen of the 264 host factors from Karlas *et al.* 2010(33) (detailed above), and in a full genome screen for influenza virus host factors in A549 cells (287/22,843 = 0.013)(33). The full genome screen provides the expected hit rate when randomly sampling the PPI. An alternative approach to network-based discovery is to target the nearest neighbors of the virus; a comparison provided by the screen of virus-interacting host proteins. An additional metric is the hit rate observed in our siRNA screen of the host factors identified by Karlas et al (71 out of 264; hit rate = 0.27). Differences between hit rates was compared using the Pearson’s chi-squared test when comparing proportions between two binomial groups.

The siRNA screen of the connecting proteins found that connecting proteins were significantly enriched for host factors, but there was no statistically significant advantage in selecting proteins by betweenness (Fig. 4). Of the 78 proteins targeted in the siRNA screen of connecting proteins, a total of 27 significantly reduced virus titers by at least two orders of magnitude; corresponding to 15 categorized as connecting – high betweenness proteins and 12 categorized as connecting – low betweenness proteins. Note that one of the 39 connecting – high betweenness proteins (PLK1) was eliminated from the calculation because both respective siRNAs were cytotoxic (see Table S4). The hit rate of connecting proteins (27/77 = 0.35) was significantly higher than the hit rate observed in the screen of virus-interacting proteins (p = 0.024) and in the full genome screen (p < 2.2*10^−16^) but not significantly distinct from the rate observed in the re-screening of the Karlas host factors (p = 0.21). When considering the connecting proteins based on their betweenness, the high betweenness had a hit rate of 0.39 (15/38) which was significantly higher than the hit rates observed in the virus-interacting and full genome screens (p= 0.032 and p < 2.2*10^−16^, respectively). High betweenness protein hit rate was higher than the rate observed in the screen of Karlas *et al.* 2010 host factors, but not significantly (p=0.16). The low betweenness connecting proteins hit rate was lower than that of the high betweenness connecting proteins (12/39 = 0.31). The difference in hit rates between high and low betweenness proteins was not significant (p = 0.57). In all, the screening results suggest that proteins connecting virus-interacting proteins to host factors of influenza virus replication are highly enriched for host factors themselves – significantly more so than proteins which directly interact with virus proteins. However, the topological information from betweenness does not significantly improve host factor identification.

**Figure 4.**
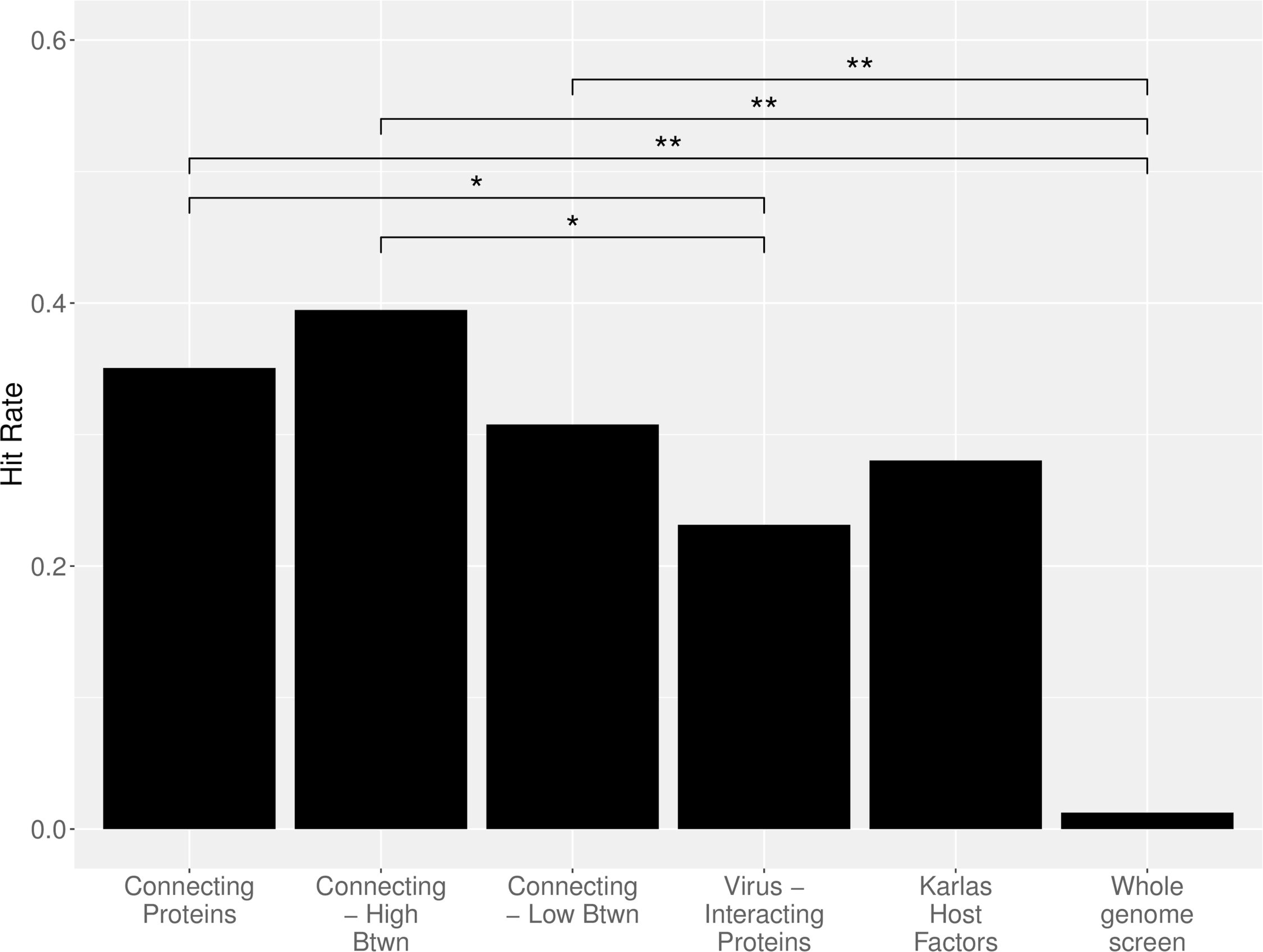
Comparison of hit rates. The hit rates are reported for all tested connecting proteins and connecting proteins with high or low betweenness in the virus subnetwork. These hit rates are compared to hit rates observed from a previous screen of virus-interacting host proteins (labeled “Virus-Interacting Proteins”) [32], from applying our screening methodology to host factors identified in a screen by Karlas *et al.* (labeled “Karlas host factors”) and from a genome-wide screen [33]. Prop.test in R was used to determine the significance of the difference in hit rates observed for binomial groups. * indicates a p < 0.05 and ** indicates a p < 0.01.

### The influenza virus subnetwork is enriched for host factors identified in 6 host factor screens

To determine if host factors identified in previous screens are enriched within the virus subnetwork, we compiled a list of host factors of influenza virus replication identified in at least one of 6 previous screens (33, 48–52) (Table S5). A Fisher exact test for enrichment was used to determine if the connecting proteins or the set of influenza virus-interacting proteins are enriched with host factors identified in these studies relative to the abundance of host factors within the PPI. Both connecting proteins and the virus interacting proteins are significantly enriched for host factors (p=7.2*10^−05^ and p=1.1*10^−05^, respectively; odds ratio = 1.4 and 1.5, respectively). When compared, there is no significant difference in the enrichment of host factors between connecting proteins and virus-interacting proteins (p = 0.48; odds ratio = 0.92). To ensure the host factors identified in the Karlas et al study were not creating bias in the enrichment result, the enrichment analysis was repeated using host factors identified in all studies except the Karlas study. Again, connecting proteins and virus interacting proteins are significantly enriched for host factors (p=1.8*10^−06^ and p=3.2*10^−03^, respectively; odds ratio = 1.5 and 1.34, respectively) and no significant difference in the enrichment of host factors between connecting proteins and virus-interacting proteins was found (p = 0.49).

## DISCUSSION

Network approaches have demonstrated their potential impact on health related research including gene/protein characterization and drug design and side effects (14, 18, 19, 22, 36, 53) yet demonstrations that network information can inform drug target discovery is still limited. Here, we present the first confirmation that virus and host protein interaction data can be integrated to improve large-scale drug target discovery (specifically antiviral target discovery) and reveal additional insights into virus-host interactions. The position of virus-interacting proteins suggest that the influenza virus has evolved to interact with proteins that heavily influence network behavior. Additionally, virus-interacting proteins are closely clustered in the network. This may be a result of attempts by the virus to manipulate specific biological functions (as proteins with shared biological functions tend to cluster in PPI networks(54)) signifying that influenza virus has parallel available pathways to engage with host biological functions. Previous studies have found that host factors of virus replication (not necessarily virus-interacting host proteins) have also been observed to cluster within the PPI network(41). Further analysis on network clustering host factors of interest is needed to determine if these two observations are independent of one another.

The observation that host-virus interaction data can be leveraged to improve virus replication host factor discovery is unlikely to be affected by off-target concerns associated with siRNA screens. Off-target concerns often challenge siRNA studies though changes to experimental protocols (such as requiring multiple siRNA hits per targeted gene or changing siRNA concentrations) can only moderately improve false positive rates (55–57). The protocol used in this study was not optimal to ensure the characterization of any one targeted gene. As such, the hit rates of gene groups are compared. Protocols between these experiments and those used for comparisons are either identical (32) or very similar (33), suggesting that off-target rates across the tested groups are unlikely to explain the differences in observed hit rates.

The variability and incompleteness of PPI data as well as the limited agreement between influenza virus replication screens are well known concerns for network-based drug target discovery. The possibility that virus-host interaction data is skewed towards well studied networks could also have an effect on the clustering result in virus-interacting proteins. However, the enrichment of host proteins important for influenza virus replication within the constructed virus subnetwork demonstrates that even with these possible shortcomings, PPI network analyses have the power to identify important host factors for influenza virus replication. The antiviral protein candidate screen performed in this study further supports the use of PPI data during candidate prioritization with significant hit rates against virus-interacting proteins and randomly targeted proteins.

The observation that betweenness does not significantly improve host factor prediction suggests that alternative topology measures should be considered. There were several reasons why betweenness was selected. Biological pathways are known to have several alternative routes to maintaining cellular operations; a key feature of biological robustness (58–60). Biological networks are also theorized to have a bow tie-like structure that suggests a natural bottlenecking within the PPI near critical cellular machinery(61). These concepts together suggest targeting bottlenecks (high betweenness proteins) as a means of mitigating escape via alternative paths. It was also the concern of alternative pathways as to why the set of virus interacting proteins was not limited to confirmed host factors of influenza virus replication. In future work, other network topology measures, e.g. degree, Burt’s Constraint, or closeness, could be tested in the subnetwork and subnetwork construction could be varied to consider different subsets of either the virus interacting proteins or the internal host factors. Even so, the results suggest that the construction of the virus-specific subnetwork provides major advantages in host factor discovery and can significantly expand drug candidate repertoires beyond virus-interacting proteins. Furthermore, since the connecting proteins do not directly interact with the virus, they may be more resistant concerns related to drug-mediated selective pressure.

Another interesting continuation of this study would identify the cause of connecting proteins’ effect on virus replication. The mechanism by which each host factor is regulating virus replication may offer additional clues for drug candidate prioritization efforts. Overall, this PPI-based study provides insight into the network characteristics of virus-host interactions and supports the idea that the influenza virus evolved to interact with host proteins in dominant network positions in order to maximally manipulate host cells.

## METHODS AND MATERIALS

### Protein-protein interaction network construction and analysis

Protein-protein interaction data was downloaded from the Human Integrated Protein-Protein Interaction rEference (HIPPIE) database(37) (version 1.4). Interactions with a confidence score less than 0.7 were removed. The interaction data was then analyzed with the igraph package in R. The interaction data resulted in one large network containing 9,969 nodes and 86 smaller disconnected networks (most with 2 nodes, all contained 7 or fewer) which were removed from the study. The final human PPI contained 9,969 proteins and 57,615 interactions.

All PPI topology analyses were performed with the R library igraph version 1.0.1(62).

### Statistical analyses and graphics packages

All statistical tests were performed in R 3.2.2 using the functions prop.test, fisher.test, pairwise.t.test or wilcoxon.test (which performs a Mann-Whitney-U test) as appropriate. Prop.test and fisher.test both compare outcome proportions between binomial groups with the latter being more precise for small group sizes. Graphics were produced with either the default graphing features of R or with the ggplot2 library (63).

### Cells and Viruses

Influenza A/WSN/ 33 virus (WSN; H1N1) was generated using reverse genetics(64). HEK293 cells were cultured in DMEM (Sigma-Aldrich) supplemented with 10% FCS (10% FCS/DMEM) and antibiotics at 37°C in 5% CO_2_. Virus plaque titers were determined by plaque assay in Madin–Darby canine kidney (MDCK) cells. MDCK cells were cultured in Eagle’s MEM (GIBCO) with 5% NCS at 37°C in 5% CO_2_.

### siRNA Treatment

siRNA treatment procedure, cell viability and virus titer determination are described in detail in Watanabe et al 2014. Briefly, two siRNAs per candidate gene were selected from a predesigned genome-wide human siRNA library (FlexTube siRNA; QIAGNE). AllStars Negative Control siRNA (QIAGEN) was served as a negative control. The siRNA against the NP gene of WSN virus (GGA UCU UAU UUC UUC GGA GUU) purchased from Sigma-Aldrich was used as a positive control. HEK293 cells were transfected twice with 25 nM (final concentration, 50 nM) of siRNA duplexes using RNAiMAX (Invitrogen). At 24 hr after the second transfection, cell viability was determined using the CellTiter-Glo assay system (Promega) following manufacturer’s instructions. To assess influenza virus replication, two parallel sets of siRNA-transfected cells were infected with 50 plaque forming units (pfu) of WSN virus per well of a 24-well tissue culture plate at 24 hr after the second siRNA transfection. At 48 hr post-infection, supernatants were harvested and virus titers determined by plaque assay in MDCK cells.

### Quantitative reverse transcription-PCR

To confirm the down-regulation of host genes by their respective target siRNAs, quantitative reverse transcription-PCR (qRT-PCR) experiments were performed. Table S6 provides a complete list of primer sequences. HEK 293 cells, transfected twice with 25 nM of siRNA (final concentration, 50 nM), were lysed at 48 h post-transfection and total RNA was extracted by using the Maxwell 16 LEV simplyRNA Tissue Kit (Promega). Reverse transcription was performed by using ReverTra Ace qPCR RT Master Mix (TOYOBO, Osaka, Japan) or SuperScript III Reverse Transcriptase (Invitrogen). The synthesized cDNA was subjected to quantitative PCR with primers specific for each gene by using the THUNDERBIRD SYBR qPCR Mix (TOYOBO). The relative mRNA expression levels of each gene were calculated by the Δ Δ Ct method using beta-actin as internal control. Primer sequences are available upon request.

### Determining candidate proteins involved in influenza virus replication

For each set of siRNAs, the virus titers from cells treated with siRNAs were normalized by the titers obtained from cell treated with AllStars Negative Control siRNA (Table S7). siRNAs that reduced cell viability by more than 40% relative to that of AllStars Negative Control siRNA-treated cells were not considered for further analysis. Unlike our previous study(32), LOESS regression was not needed (Fig. S3). A two-sided, unpaired Student’s t test was used to compare the mean virus titers in cells treated with gene-specific siRNAs with those in cells treated with AllStars Negative Control siRNA. Holm’s method for multiple comparisons was then applied to the p values.

## ACKNOWLEDGEMENTS

We thank Naomi Fujimoto, Tomoko Kuwahara and Kazue Goto for technical assistance.

## DECLARATION OF INTEREST

The authors declare no competing interests.

## SUPPLEMENTAL FIGURES

**Fig S1 The distributions of the (a) degree and (b) betweenness of the interaction partners of each of the 11 virus proteins.** The y axis lists the particular virus protein, and the x axis demonstrates distributions of the centrality measures of the virus protein’s interaction partners within the human PPI network. The distributions for all proteins in the human PPI network (labeled “All”) and the set of proteins that interacted with any of the virus proteins (“VB”) are included for comparison.

**Fig S2 Boxplot of the degree and betweenness distributions for connecting (candidate) proteins, virus-interacting proteins, and internal essential proteins.** Black lines indicate the median for each population.

**Fig S3 The mean log fold change (LFC) vs the mean fold change (FC) in cell viability for all 156 gene-specific siRNAs tested.** Cyan and green points highlight data corresponding to the 24 negative and positive control siRNAs (i.e., AllStars Negative Control siRNA and 25 siRNA against influenza virus NP gene, respectively). The broken ride line is the LOESS regression curve, showing that virus growth was not dependent on cell viability.

**Table S1 Effects of siRNAs targeting host factors identified to be important for influenza virus replication by Karlas et al. (Nature, 2010) on virus production.** Note that two siRNA’s were used per Entrez Gene ID. Sheet 2, labeled “untested host factors”, lists host factors that were identified in the Karlas screen but were not evaluated in this study.

**Table S2 The degree and betweenness of proteins in virus-host interaction subnetwork.** The symbol, description and Entrez Gene ID of each protein are provided in the first three columns. Proteins tagged with a 1 in the “Virus-interacting” and “Internal-Essential” columns identify proteins were associated with a virus protein in the co-immunoprecipitation study or identified as essential but not directly associated with a virus protein, respectively. The last three columns provide the protein’s degree and betweenness in the subnetwork and identify which proteins were selected for further testing.

**Table S3 DAVID Functional Annotation Tool results for virus-interacting proteins and connecting proteins of the influenza virus subnetwork.** Full results include the clustering, chart, and table outputs from DAVID 6.8.

**Table S4 Effects of siRNAs Targeting Host Factors with High or Low Betweenness in the Virus-Host Subnetwork on Virus Production.**

**Table S5 Hit-lists of genes identified in 6 independent genome-wide screens.** Studies include König et al (2010), Brass et al (2009), Shapira et al (2009), Hao et al (2008), Karlas et al (2010), and Sui et al (2009).

**Table S6 A list of primers used for qPCR.**

**Table S7 Virus titers observed in HEK293 cells.**

